# Data Driven Taxonomy for Antipsychotic Medication: A New Classification System

**DOI:** 10.1101/2023.01.23.524854

**Authors:** Robert A. McCutcheon, Paul J. Harrison, Oliver D. Howes, Philip K. McGuire, David Taylor, Toby Pillinger

## Abstract

**Background:** There are over 25 licensed antipsychotic medications with diverse pharmacological and clinical profiles. Antipsychotics are commonly described as either ‘typical’ or ‘atypical’, but this does not accurately reflect pharmacological profiles. There is thus a need for a data driven antipsychotic classification scheme suitable for clinicians and researchers which maps onto both pharmacological and clinical effects.

**Method:** We analysed affinities of 27 antipsychotics for 42 receptors from 3,325 receptor binding studies. We used a clustering algorithm to group antipsychotics based on their pattern of receptor affinity. Using a machine learning model, we examined the ability of this grouping to predict antipsychotic-induced side effects quantified according to an umbrella review of clinical trial and treatment guideline data.

**Results:** Clustering resulted in four groups of antipsychotics. The predominant receptor affinity and effect/side effect ‘fingerprints’ of these four groups were defined, as follows:

Group 1 - Muscarinic (M3-M5) receptor antagonism; Cholinergic and metabolic side effects.

Group 2 - Dopamine (D2) partial agonism and adrenergic antagonism; Globally low side effect burden.

Group 3 - Serotonergic and dopaminergic antagonism; Globally moderate side effect burden.

Group 4 - Dopaminergic antagonism; Extrapyramidal and motor side effects.

Groups 1 and 4 were more efficacious than clusters 2 and 3. The novel classification was superior to existing approaches when predicting side effects.

**Conclusions:** A receptor affinity-based grouping not only reflects compound pharmacology but also detects meaningful clinical differences to a greater extent than existing approaches. The approach has the potential to benefit both patients and researchers by guiding treatment and informing drug development.

## Introduction

Psychotropic agents have traditionally been classified based on clinical indication.(1) In the case of ‘antipsychotics’, the drugs used to treat schizophrenia and related psychoses, this classification has also mapped onto a shared pharmacological mechanism of D2 dopamine receptor antagonism, which is tightly linked to clinical efficacy.(2–4)

Despite sharing a common dopaminergic mechanism of action, there are significant differences between antipsychotic agents both in terms of their broader pharmacological and clinical effects.(5–7) Early attempts to provide a more granular classification of antipsychotics have employed an ‘atypical/typical’ or (almost identical) ‘first/second generation’ dichotomy.(8) While initially proposed to reflect mechanistic differences it has subsequently become clear that the compounds within these categories shared neither common pharmacological or clinical profiles.(9) The subsequent development of a ‘Neuroscience based Nomenclature’ (NbN) was motivated in part to address this shortcoming.(9,10) The NbN approach categorises compounds by clinical indication and a summarised receptor profile. However, this process relies to some reliant on expert judgment and involves a simplification of the highly diverse pharmacology of this group of compounds. Although simplification may be necessary when developing a system that can be applied across the pharmacopeia, it has the potential to obscure important similarities and differences between drugs.

There are large interindividual differences in antipsychotic response and many patients switch antipsychotics multiple times before finding one that is both well tolerated and effective.(11–13) There is currently no grouping of antipsychotics to guide clinicians and patients in their choice of initial or subsequent drug. For patients whose psychotic symptoms have not improved adequately with first-line treatment or who are experiencing side-effects, clinical guidelines recommend switching to a different antipsychotic but give little guidance on which drug to select.(14,15) Even where there is guidance, this is typically limited to switching between atypical/typical agents, which does not reflect efficacy or side-effect profiles.(14) Thus, a classification system that facilitates a switch to a second-line agent with a distinct pharmacological mechanism of action may improve chances of treatment response and/or tolerability. Recognising that antipsychotics with similar receptor binding profiles share similar side effect profiles may also help drug development. For example, dopamine receptor partial agonists have been heralded for their relatively more benign metabolic side effect profiles;(16) however, in a recent network meta-analysis ranking antipsychotics based on their associated metabolic side effects, ziprasidone and its structural analogue lurasidone (both dopamine receptor antagonists) were grouped with the partial agonists as superior agents.(5) Comprehensively understanding patterns of pharmacological similarity across compounds may support initiatives to develop safer and more tolerable treatments.

A systematic synthesis of the pharmacology of antipsychotic medication is made possible by the availability of a high number of receptor binding studies covering a wide range of receptor types.(17) These studies enable the construction of a receptor ‘fingerprint’ for individual antipsychotics. In the current paper, we synthesise the results of all relevant receptor binding studies to derive a receptor fingerprint for each antipsychotic. We then apply an unbiased clustering algorithm to group antipsychotics with similar profiles, before developing a machine learning model that uses receptor profiles to predict efficacy and side effect burden. We find that receptor-defined groupings show limited overlap with existing classification schemes but a good mapping to clinical profile, and by definition to receptor profiles.

## Methods and Materials

### Overview

We performed a comprehensive search for antipsychotic receptor affinities. We then clustered antipsychotics into groups based on the similarity of their receptor profiles. We next characterised these receptor-defined clusters in terms of their receptor affinities and clinical profiles. Finally, we compared the ability of these clusters to predict side effects and compared this to existing methods of categorising antipsychotics.

### Determining receptor affinities

As in previous work,(18) receptor affinities for all licensed antipsychotic drugs were obtained from the National Institute of Mental Health Psychoactive Drug Screening Program (PDSP) database https://pdsp.unc.edu/databases/kiDownload/.(17) Only data from studies that reported binding to human tissue were included. A receptor was included in the analysis if data were available for at least 5 separate drugs. Antipsychotic drugs were included in the subsequent analysis if data were available for at least 5 separate receptors. Receptors were removed if Ki values were identical for all drugs. If multiple studies existed for the same receptor and drug, then the median value was calculated and used in subsequent analyses. Finally, Ki values were converted to pKi values.

### Clustering antipsychotics based on receptor affinities

The pKi values were first reduced by 4 as this produced a floor score of zero. Next, in the case that a drug was an agonist or partial agonist at a given receptor, the pKi value for that drug-receptor combination was multiplied by -1 to account for the functionally inverse effect. Without this inversion there would be no distinction between agonists and antagonists.

Probabilistic principal components analysis (PPCA) was then used to impute any missing pKi values.(19) Then, the adjusted pKi values for all antipsychotics were Pearson correlated with one another to produce a correlation matrix. In this correlation matrix a high correlation coefficient between two antipsychotics indicates that they share a similar receptor profile. This approach has a similar effect to normalising for D2 pKi,(18) thereby accounting for the dosing differences between antipsychotics. The Louvain clustering algorithm was then used to group antipsychotics with similar receptor profiles into distinct groups.(20)

### Characterising the relationship between receptor profiles, categorisation schemes, and clinical effects

To characterise the receptor profile of the antipsychotic clusters identified above we performed a PPCA of the receptor profiles, then for each cluster calculated the mean component loading for the three components explaining the greatest proportion of variance.

Relative side effect burden (magnitude or relative risk) for 13 common adverse effects (weight gain, Parkinsonism, akathisia, anticholinergic effects, sedation, hyperprolactinaemia, QT prolongation, orthostatic hypotension, dystonia, tardive dyskinesia, seizure, dyslipidaemia, and dysglycaemia) and efficacy (in terms of positive, negative, and total symptoms) of antipsychotics included in the PPCA analysis were obtained from an umbrella review of network meta-analyses and clinical guidelines for the acute treatment of schizophrenia (see supplementary). For each side effect and efficacy measure, we characterised the mean for each of the 4 receptor-defined clusters.

We next examined whether complete receptor binding profiles, and receptor-profile based groupings (clusters), were predictive of clinical profiles, and compared with existing classification schemes. We developed a prediction model using training data consisting of all but one of the available antipsychotics. Within the training data any missing side effect values were imputed using PPCA. In this model either the receptor profiles (number of predictor variables, D=42), the receptor profile defined groupings (D=4), NbN defined groupings (D=7), or a typical/atypical/partial agonist grouping (D=3) was used as the predictor variables, while the side effect and efficacy scores (k=16) were used as the target variables. The NbN groupings were defined on the basis of their mode of action as reported at https://nbn2r.com/authors. A definitive typical/atypical distinction is not available, with most drugs (other than clozapine) classified according to year of discovery, the classification adopted for the current analysis is displayed in Figure 5.

**Figure 1.**
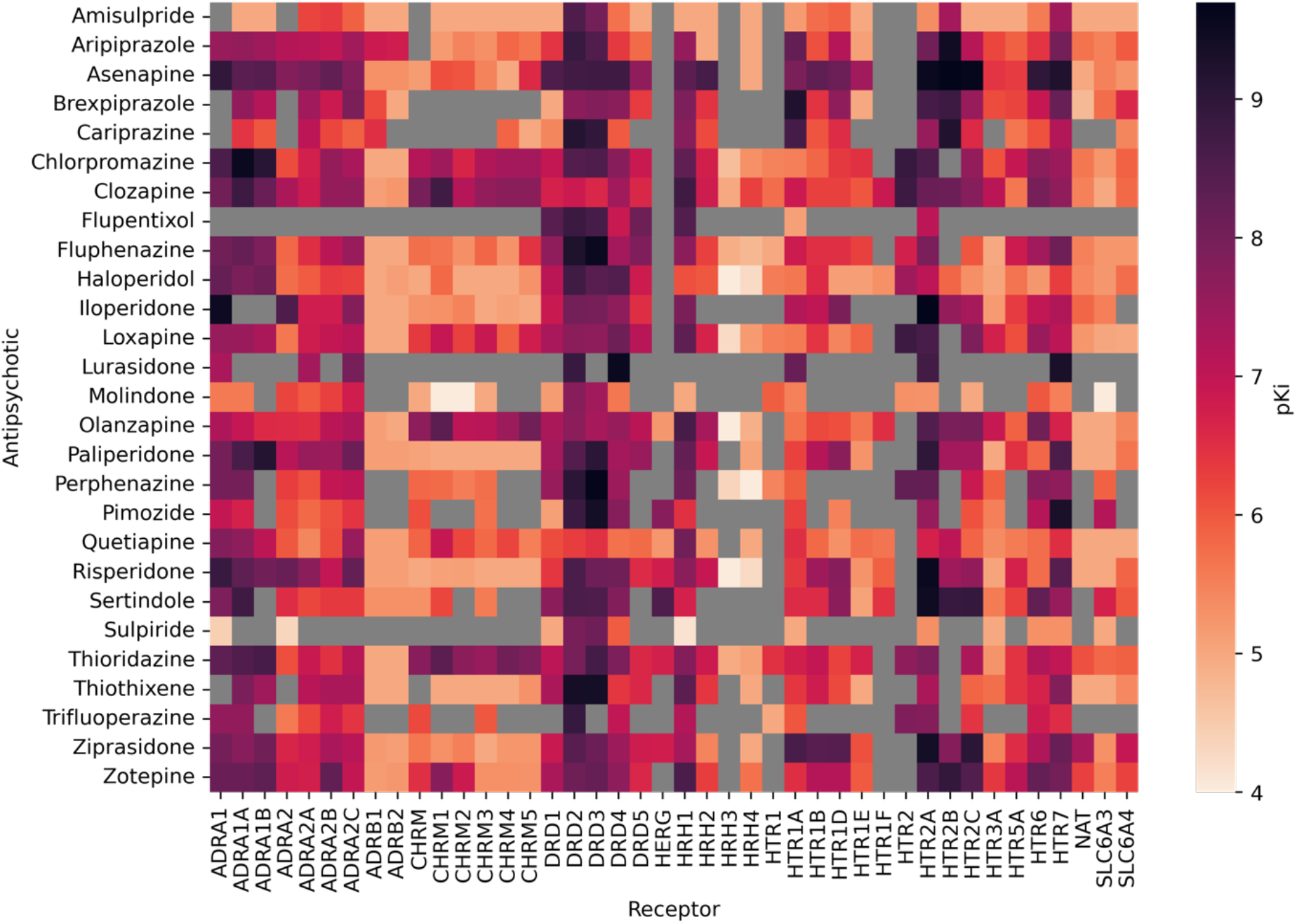
Antipsychotic pKi values. A larger pKi indicate greater affinity of the drug to receptor. For visualisation purposes data here represents pKi values with no adjustments made on the basis of whether a drug is an agonist or antagonist, whereas subsequent analyses make this adjustement. Gray square indicate an absence of data. ADRA: Alpha adrenergic receptor, ADRB: Beta adrenergic receptor, CHRM: Muscarinic acetylcholine receptor, DR: Dopamine receptor, HERG: Human ether-a-go-go-related gene, HR: Histamine receptor, HTR: Serotonin receptor, NAT: Noradrenaline transporter, SLC6: Solute carrier family 6 transporter

**Figure 2.**
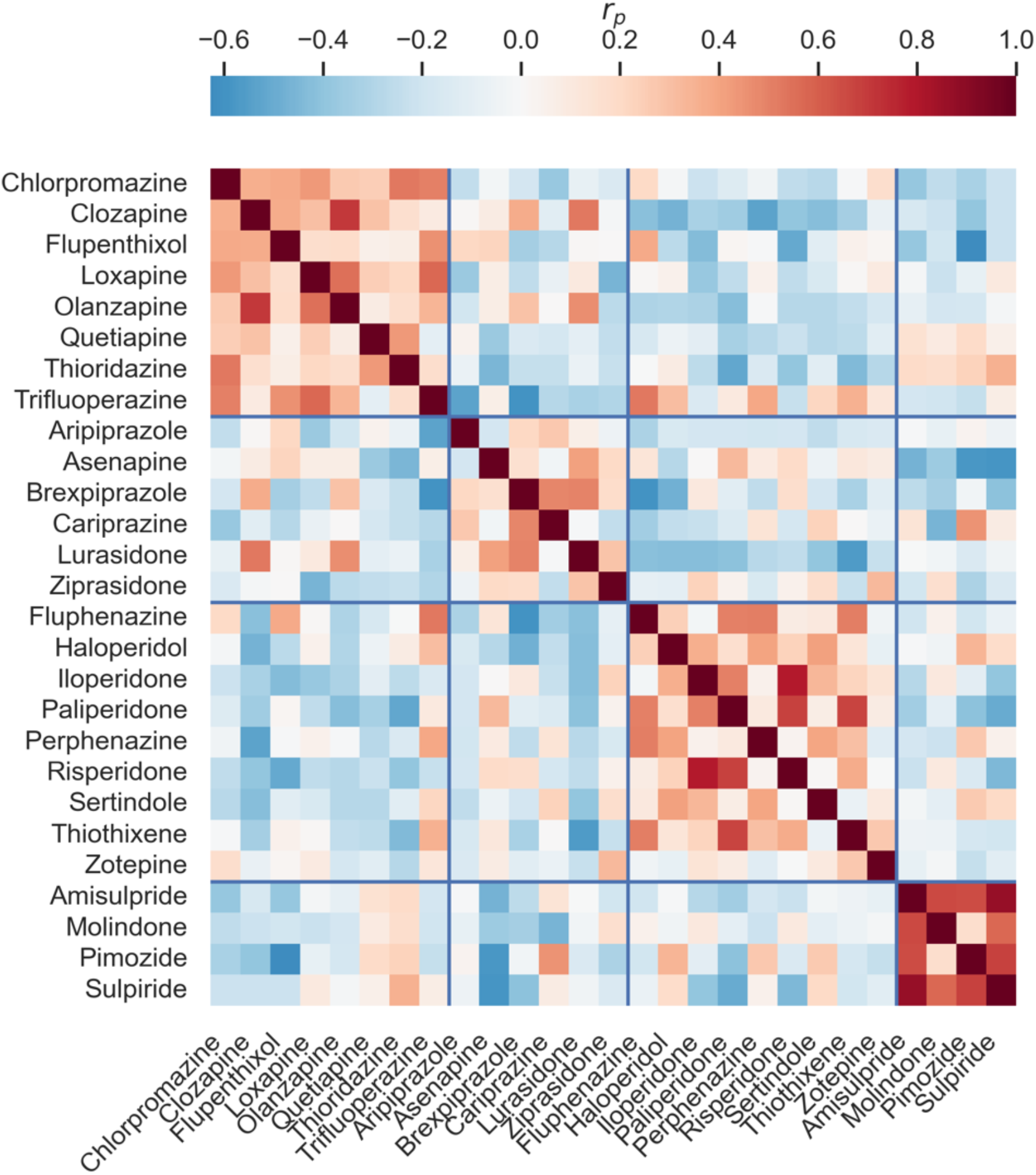
Antipsychotic clustering based on receptor profiles. The colour of each small square indicates the strength of correlation between the receptor profile of the antipsychotic in the corresponding row and column (e.g. one can see that pimozide shows a similar receptor profile to amisulpride but not to flupentixol). The grouping outlines by the blue lines reflects the result of a clustering algorithm that aims to group highly correlated drugs together.

**Figure 3.**
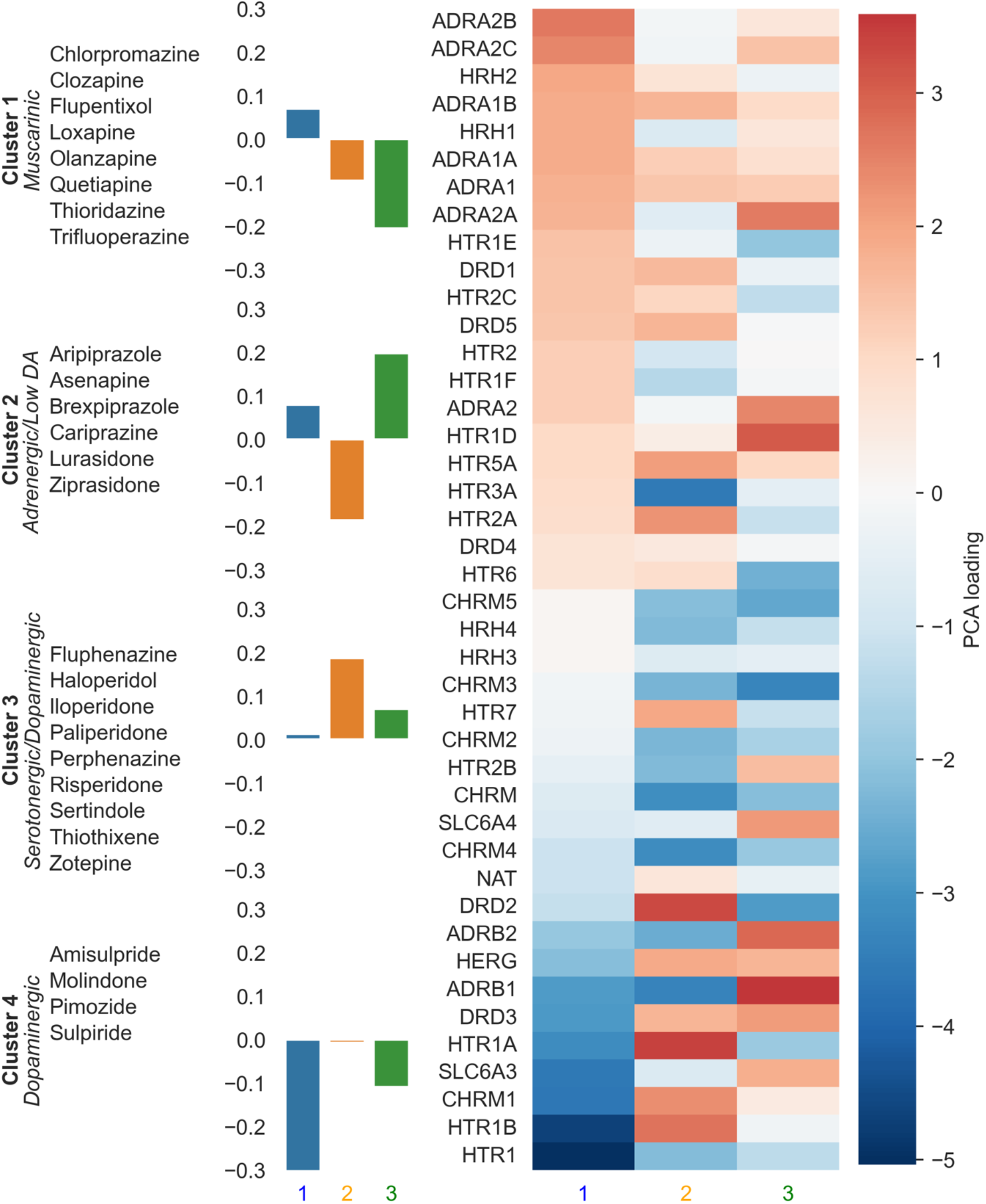
Characterising receptor defined antipsychotic clusters. The numbers ‘1’, ’2’, and ’3’ refer to the first three principal components The bar chart shows that e.g. cluster 4 has a large negative loading for the component 1. The heatmap shows how the components relate to the receptor profile. The large negative loading for component 1 in cluster 4 indicates that the drugs in this cluster will tend to act as relatively strong antagonists at HTR1 and CHRM1, and weak antagonists (or even agonists) at ADRA2B, and ADRA2C.

**Figure 4.**
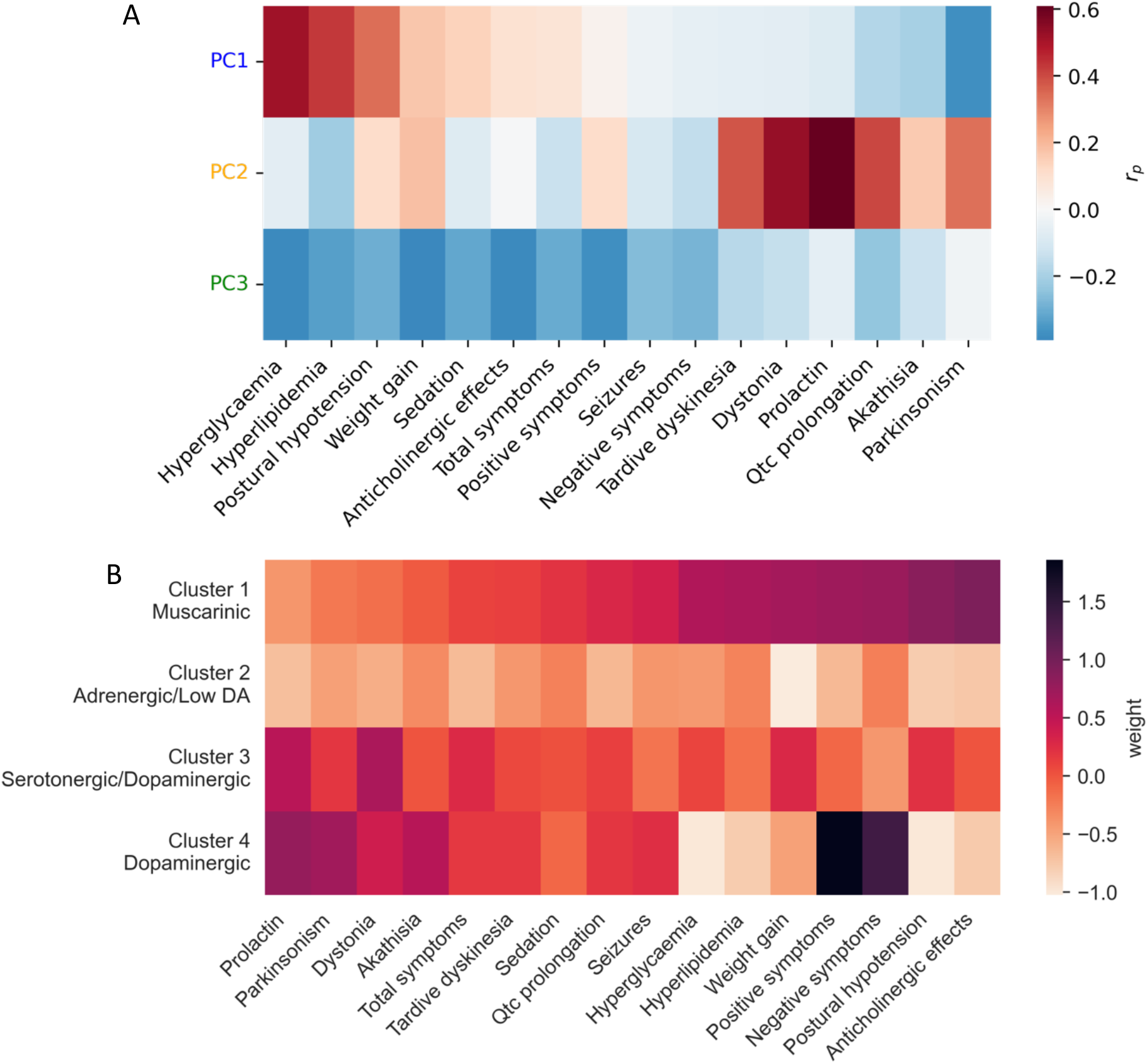
Characterising clinical profiles of principal components and receptor defined clusters. A) Correlation coefficients across antipsychotics between principal component loadings illustrated in Fig 3 and side effect and efficacy scores. Red indicates that a drug with a high loading for that component is likely to be associated with the effect in question. B) Mean scores for antipsychotic clusters illustrated in Figure 2, a darker colour indicates that cluster is associated with greater severity of the side effect in question.

**Figure 5.**
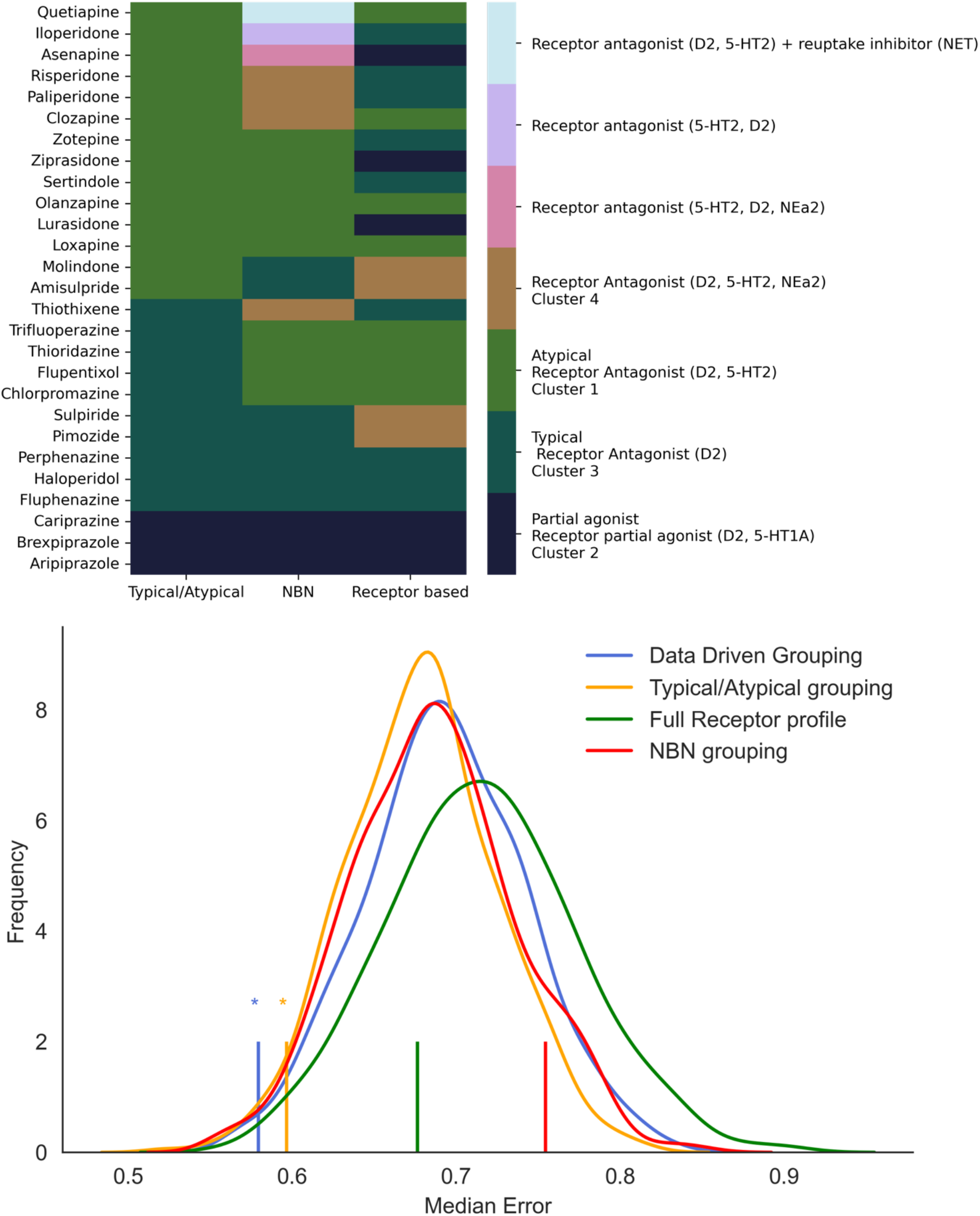
Antipsychotic categorisation schemes and prediction of clinical effects. (A) Antipsychotics classified according to a typical/atypical/partial agonist split, Neuroscience based Nomenclature (NBN), and the receptor defined clusters illustrated in Figure 2. (B) The curves illustrate the permutation generated null distribution . Vertical lines indicated the observed median error for predicting out of sample side effect profiles (a smaller value reflects more accurate prediction). The data driven and typical/atypical groupings produce a statistically significant prediction of overall clinical profile compared to the null distribution.

We used partial least squares regression, given this is a model well suited to using multiple features to simultaneously predict multiple targets. We then used the partial least square model that had been fitted on the training data to predict side effects for the single antipsychotic not included in the training data. We then calculated the median of the absolute error between predicted and observed side effect scores for this antipsychotic. We repeated this across all 27 antipsychotics and calculated the median error score across the 27 antipsychotics to provide a summary estimate of predictive ability for each of the four methods of categorisation. We used permutation testing to assess statistical significance of the prediction by comparing the observed median error score to a null distribution of median error scores generated by shuffling antipsychotics in the training data 500 times so as to break the connection between receptor and side-effect profile. Code and data for all analyses is available at https://github.com/rob-mccutcheon/antipsychotic_pca_paper

## Results

### Receptor affinities

In total, 97,599 Ki values were extracted. Of these, 5,304 related to antipsychotics, of which 3,325 reported on binding to human tissue. Data regarding 67 distinct receptors and 29 different antipsychotics were reported but this was reduced to 42 receptors and 27 antipsychotics when the requirement for >5 datapoints for each receptor and drug were applied (2 drugs and 13 receptors were removed, see supplementary for details). The pKi values are displayed in figure 1.

### Clustering antipsychotics based on receptor affinities

Antipsychotics were clustered based on the similarity of their receptor affinity profiles. Four clusters were identified, and are illustrated in Figure 2.

To summarise the receptor affinity profile of each cluster, we examined the mean loading for the three PPCA components explaining the greatest variance (see Figure 3). The first cluster (chlorpromazine, clozapine, flupenthixol, loxapine, olanzapine, quetiapine, thiordiazine, and trifluoperazine) was characterised as ‘muscarinic’ given its strong negative loading on the third component which reflected antagonism at the muscarinic M3-M5 receptors (but either agonism or weak antagonism at the M1 receptor). The second cluster (aripiprazole, asenapine, brexpiprazole, cariprazine, lurasidone, and ziprasidone) was characterised as ‘adrenergic with low dopaminergic antagonism’ had a strong positive loading on the third component and a strong negative loading on the second component reflecting a lack of muscarinic or serotonergic antagonism but significant adrenergic antagonism, and dopamine D2 partial agonism. The third cluster (fluphenazine, haloperidol, iloperidone, paliperidone, perphenazine, risperidone, sertindole, thiothixene, and zotepine) was characterised as ‘serotonergic-dopaminergic’ due to its strong positive loading on the second component which reflects serotonergic and dopaminergic antagonism. The fourth cluster (amisulpride, molindone, pimozide, and sulpiride) was characterised as ‘dopaminergic’ given its strong negative loading on the first component, which reflects relatively pure dopaminergic antagonism without adrenergic effects.

We then characterised how these principal components of receptor affinity correlated with efficacy measures propensity to cause side effects (see Figure 4A). Drugs with a high loading for the first principal component are more likely to cause metabolic and cholinergic side effects, but a low propensity for Parkinsonism, akathisia, and hyperprolactinaemia, they also show the greatest efficacy for total symptoms. Drugs with a high loading for the second component show the opposite pattern, with a relative propensity to cause Parkinsonism, akathisia, and hyperprolactinaemia over metabolic effects, they also show the greatest efficacy for positive symptoms. A high loading for the third component reflects a propensity for a generally low all round side effect burden but also less efficacy in terms of total, positive, and negative symptoms.

We also looked at the mean side effect levels for the receptor defined clusters (see Figure 4B). We found that cluster 1 was associated with anticholinergic side effects, postural hypotension, and metabolic side effects, cluster 2 was associated with a globally low side effect burden, cluster 3 with a globally moderate burden, and cluster 4 with parkinsonism, akathisia and hyperprolactinaemia. Clusters 1 and 4 were more efficacious than clusters 2 and 3.

Finally, we examined which of 4 classification methods (the receptor profile defined grouping described here, complete receptor binding profiles, NbN defined grouping, and atypical/typical/partial agonist defined groupings, see Figure 5a) were able to best predict out of sample side effect profiles (see Figure 5b). We found that both receptor defined clusters described in the current paper (p=0.01) and typical/atypical/partial agonist grouping (p=0.04) produced statistically significant predictions, in contrast to complete receptor profiles (p = 0.25), or NbN grouping (p = 0.90).

## Discussion

This paper illustrates how receptor profiles can be used to classify antipsychotics in a data-driven fashion. Furthermore, we demonstrate that the groupings derived from this approach predict side effect profiles with greater accuracy than existing classification schemes. The receptor defined grouping even performed better than the entire receptor profile in predicting clinical profiles, likely reflecting overfitting given the relatively low number of samples in comparison to number of receptors included.

These findings have several implications. An unbiased pharmacologically driven approach to classification has a priori advantages in that, by definition, it reflects pharmacology and does not require decisions as to which receptors to prioritise. In addition, we have demonstrated how receptor profiles can be used to quantitatively estimate side effects, which has potential uses when evaluating compounds that have not yet undergone clinical testing. Furthermore, treatment side effects are key factors that people with schizophrenia and their clinicians consider when making prescription decisions,(21) and reducing side effects of antipsychotics is central to initiatives to improve morbidity and mortality rates in this patient group.(22) Although treatment decisions based on side effect burden may be best made at the individual drug level, we have identified groups of antipsychotics with similar receptor binding signatures and either globally low or moderate side effect burdens; this has the potential to inform clinical practice. For example, we identified a group of antipsychotics with a low side effect risk that included all licensed partial agonists alongside ziprasidone and its structural analogue lurasidone. Previous studies and clinical guidance documents have recommended that this same group of antipsychotics be selected preferentially when there is a desire to avoid metabolic side effects;(5,16) this is consistent with our data-informed classification scheme but our scheme extends this to indicate that they are preferential in terms of other side-effects as well. Thus, guidelines and clinicians may recommend a drug from this group as first line treatment given the overall favourable side-effect profile and as a rational choice to switch to for patients experiencing metabolic side-effects on a drug not in this class. In contrast, if efficacy is paramount then it may be preferable to consider clusters 1 or 4, with the decision between these dependent on whether movement or metabolic side effects are a greater concern.

In terms of treatment effectiveness, it is unclear how to select a second antipsychotic in the case of initial non-response. There is evidence that switching to a pharmacologically distinct compound produces clinical benefits.(13) Although current guidelines do recommend switching to a different antipsychotic class prior to considering clozapine, this guidance is often limited to switching between atypical/typical agents.(14) The current classification separates drugs into classes as pharmacologically distinct from one another as possible, potentially providing some guidance as to sensible switching choices when changing medication secondary to lack of effectiveness. While antipsychotic polypharmacy should typically be avoided, the classification could also be of use in suggesting more effective antipsychotic combination strategies where other options have been exhausted.(23,24) Further work definitively testing whether these groupings reflect an optimal switching strategy is warranted.

Despite these advantages, alternative taxonomies have benefits in other aspects. For example, the NbN approach encompasses psychopharmacological treatments as a whole, as opposed to solely antipsychotics. Future work, however, could similarly extend the current approach to a wider range of compounds. A drawback of the NbN approach is the necessary limitation to covering an incomplete range of receptor systems. While this has the benefit of keeping the number of systems at a manageable number for the user, it means that important facets are neglected. For example, histaminergic affinities do not feature in the NbN approach to classifying antipsychotics.(25) This is a significant limitation of the NbN approach given the central role of the histaminergic system in determining the propensity of a drug to induce weight gain.(25,26) The fact that histaminergic affinities contribute to the clustering approach employed in the current analysis may therefore be one of the factors that improve its ability to predict side effect profiles compared to NbN.

In the current analysis we attempt to move away from pre-existing biases by employing an agnostic classification algorithm. However, to obtain more fully unbiased results one requires unbiased data in addition to an unbiased algorithm. The database chosen does not reflect an entirely systematic survey of receptor affinities but to an extent reflects research interests. This means that for some drugs, such as lurasidone, flupentixol, and sulpiride, there is a paucity of data, although for the rest the database is relatively comprehensive. However, whilst this is a potential limitation of our approach, it is even more of a potential limitation for the NbN approach as it uses a sub-set of known receptor affinities. Moreover, it is partially mitigated by the fact that drugs typically undergo standard receptor affinity screening, and an advantage of our approach of alternatives is it can be readily updated with new findings as they emerge (we have made the code required openly available).

One of the main areas where our methods could potentially be improved is to account for additional pharmacological properties. Our approach focuses on relative receptor affinities, and therefore while this adjusts for dosing differences, it does not consider that active metabolites may have quite different pharmacodynamic effects to the parent compound. The fact that compounds differ in their ability to cross the broad brain barrier is also not accounted for, this means the relationship between lower permeability and greater peripherally mediated side effects will not be reflected. Finally, in the cases where individual receptors mediate the majority of an effect (e.g. HERG and QTc prolongation) it may be that the importance of the receptor becomes lost in the analysis.

Future work may consider a data driven approach to clustering all neuropsychiatric medication as opposed to solely medications used in the treatment of psychosis. To build upon the current approach, clinical studies could evaluate the benefits of the classification scheme in guiding switching decisions, while the predictive model may be of use in identifying ideal receptor profiles that maximise efficacy while minimising side effects.

In conclusion, the current study provides a pharmacological data driven approach to the classification of antipsychotic medication. We derive four groups of antipsychotics with distinct receptor, efficacy, and side effect profiles. This approach reflects the pharmacological properties as closely as possible and also shows considerable mapping to side effect profiles, suggesting that it may hold some advantages over existing approaches. This promises to benefit both patients and researchers, guiding appropriate treatment and future drug development.

## Acknowledgements

RAMs work is funded by a Wellcome Clinical Research Career Development Fellowship (224625/Z/21/Z).

TPs work is funded by an NIHR Academic Clinical Lectureship.

We thank Professor Philip Cowen for his comments on the manuscript.

## Conflicts of Interest

Dr McCutcheon has participated in advisory/speaker meetings organised by Otsuka, Karuna and Janssen. ODH is a part-time employee of H. Lundbeck A/S and has received investigator-initiated research funding from and/or participated in advisory/speaker meetings organised by Angellini, Autifony, Biogen, Boehringer-Ingelheim, Eli Lilly, Heptares, Global Medical Education, Invicro, Jansenn, Lundbeck, Neurocrine, Otsuka, Sunovion, Rand, Recordati, Roche and Viatris/ Mylan. Dr Howes has a patent for the use of dopaminergic imaging. David Taylor reports grants and personal fees from Janssen, Sunovion, Recordati and Mylan, and personal fees from Accord, outside the submitted work. Dr Pillinger has participated in educational speaker meetings organised by Lundbeck, Otsuka, Sunovion, Janssen, Schwabe Pharma and Recordati. The other authors declare no conflicates of interest.

