## Supplement for "Data Driven Taxonomy for Antipsychotic Medication: A New Classification System"

**Methods**

*Clustering*

Prior to probabilistic PCA pKi values were standardised by subtraction of the mean and scaling to unit variance. The Louvain clustering algorithm was used to group antipsychotics with similar receptor profiles. The gamma parameter was set at ‘1’, negative and positive correlations were equally weighted, and the algorithm was ran 100 times with the solution showing the greatest modularity chosen.

*Side effects data*

We performed a systematic ‘umbrella’ review of meta-analyses to create a database of effect size magnitudes for antipsychotic side effects. As part of another ongoing project we simultaneously searched for antidepressant-associated effects. We searched Pubmed from inception to August week 4 2022. We focused on key side effects described in The Maudsley Prescribing Guidelines in Psychiatry.^1^ As such, the following search terms were used: [(antidepressant OR antipsychotic) AND (extrapyramidal OR parkinsonism OR dyskinesia OR dystonia OR akathisia OR prolactin OR headache OR agitation OR insomnia OR cholinergic OR gastrointestinal OR constipation OR nausea OR arrythmia OR QTc OR hypotension OR hypertension OR weight OR glucose OR lipid OR cholesterol OR triglyceride OR sedation OR natraemia OR natremia OR sodium OR thromboembolism OR sexual) AND “meta-analysis”]. Only meta-analyses based on data from randomised controlled trials were considered. Where there was more than one meta-analysis for a given side effect, the study with the largest sample size was selected. For each side effect and medication, we extracted effect size magnitude compared to placebo. Where fewer than 2/3 of antidepressants or antipsychotics had meta-analytic side effect estimates available, we aimed to use more comprehensive side effect data sources in the form of ordinal rankings from national/international guidelines or consensus statements for treatment of depression^1–11^ and schizophrenia.^1,12–18^ These guidelines were selected in line with previous meta-reviews in the field.^19^ The resultant side effect database consisted of rows corresponding to drugs, and columns corresponding to side effects, with entries corresponding to either meta-analysis derived effect sizes or guideline-defined ordinal scores. Each side effect was then normalised (minimum-maximum scaled by subtracting the minimum value for that side effect and dividing by the range) to give values between 0 and 1.

Of 2060 citations retrieved, 11 meta-analyses met inclusion criteria. For antipsychotics, side effect data were available for 32 drugs. Six side effects (weight gain, Parkinsonism, akathisia, anticholinergic effects, sedation, and hyperprolactinaemia) had meta-analytic data available for ≥66% of available drugs (≥22 drugs); these data were extracted from 2 NMAs.^20,21^ Antipsychotic side effect data for remaining side effects were derived from national/international guidelines or consensus statements. Up to 4 guidelines/consensus statements^1,16–18^ provided ordinal rankings of 7 antipsychotic-related side effects for which sufficient meta-analytic data were not available (namely: QT prolongation, orthostatic hypotension, dystonia, tardive dyskinesia, seizure, dyslipidaemia, and dysglycaemia). Ordinal scores for drug side effects were extracted from each guideline and normalised (minimum-maximum scaled by subtracting the minimum value for that side effect and dividing by the range); where there were multiple guidelines, a mean of normalised scores was calculated. Missing side effect values were imputed using probabilistic PCA

*Partial Least Squares*

Side effect scores were highly skewed, therefore scores were shifted to have a zero floor before undergoing a square root transform and standard scaling (removal of mean and scaling to unit variance). Scaling was fit only on training data, before later being applied to left out test sample. Partial least squares was implemented using the nipals algorithm with 3 components.
